# The Role of Similarity of Stimuli and Responses in Learning by Nectar-Foraging Bumble Bees: A Test of Osgood’s Model

**DOI:** 10.1101/2024.04.25.591145

**Authors:** Minjung Baek, Daniel R. Papaj

## Abstract

Learning stimulus – response associations helps animals to adjust to changing environments. Sequentially learned associations may interact with each other, either reinforcing memory, a process referred to as ‘transfer’, or hindering memory, a process referred to as ‘interference.’ According to Osgood’s (1949) model, close similarity between new and previously learned stimuli can enhance the transfer of memory through a process of stimulus generalization. In contrast, the model proposes that if responses are different from those previously learned, generalizing stimuli may lead to confusion, resulting in the interference of memory. Except for some work in humans, the interaction between stimulus similarity and response similarity is poorly documented. Here, we tested Osgood’s model using bumble bees (*Bombus impatiens*) foraging for sucrose on artificial flowers with varied colours (= stimuli) that required either legitimate visitation or nectar robbing (= responses). Bees were first allowed to forage on one type of flower, were then switched to another, and finally were returned to the initial flower type. We measured learning performance via flower handling time and the number of failed visits. Consistent with Osgood’s model, bees made more failed visits when they switched between similarly coloured flowers requiring different foraging techniques but made fewer failed visits when switching between similarly coloured flowers with the same technique. Regardless of similarities in stimuli or responses, however, experienced bees were faster in handling flowers than were naïve bees. Results taken together thus provided mixed support for Osgood’s model. Possible explanations for the mixed results are discussed.

## Introduction

Animals must learn to behave appropriately in diverse situations, which involves formation of multiple memories over time. These multiple memories can either strengthen or weaken each other. Enhancement of memory is termed transfer, while the hindrance of memory is termed interference (Adams, 1987; Anderson, 2005; Osgood, 1949). Interference from competing memories has been identified as an important factor in forgetting in humans and animals (Anderson, 2003; Davis & Zhong, 2017). Transfer and interference of memory can act in two ways: proactively, where previously-established memory affects formation of new memory, or retroactively, where newly-formed memory affects previously-established memory. The extent to which memory transfer or interference occurs depends in part on how similar the old and new memories are. For instance, learning a task might be easier if it resembles a familiar task (Hughes & O’Brien, 2001). In this case, response similarity aids transfer. However, similar memories can also compete more strongly, leading to confusion in tasks, an example of similarity-aided interference (Lewis & Kamil, 2006; Roberts, 1981).

These seemingly contradictory effects of similarity on memory have been reported in semantic studies, where human participants were assigned the task of learning word pairs, consisting of a stimulus word and a response word (Anderson & Neely, 1996; Antony et al., 2022; Osgood, 1949). To resolve this contradiction, Osgood (1949) proposed over 70 years ago that both the stimulus similarity and the response similarity in stimulus – response associations should be considered in predicting transfer or interference of sequentially acquired memory. His transfer surface model (Figure 1a) suggests that transfer and interference are part of the same process but acting in opposite directions. The model highlights three possible effects of one learned association on another. First, when stimuli vary and responses are identical, transfer of memory will occur. The degree of transfer increases as the novel stimulus becomes more similar to the previously learned stimulus, resulting in stimulus generalization (green line in Figure 1a). Second, when responses differ, interference may occur depending on how similar stimuli are. Close similarity of stimuli promotes generalization, which leads to incorrect transfer of the previously learned response to the novel stimulus (red line in Figure 1a). Third, when stimuli are identical, transfer is maximal if responses are identical, and decreases as responses become less similar. If responses are different enough, interference results, with interference increasing in magnitude as responses become less similar (blue line in Figure 1a). Elements of the Osgood’s model have found support from empirical results, but the entire model has not been thoroughly evaluated until recently in humans (Antony et al., 2022).

**Figure 1.**
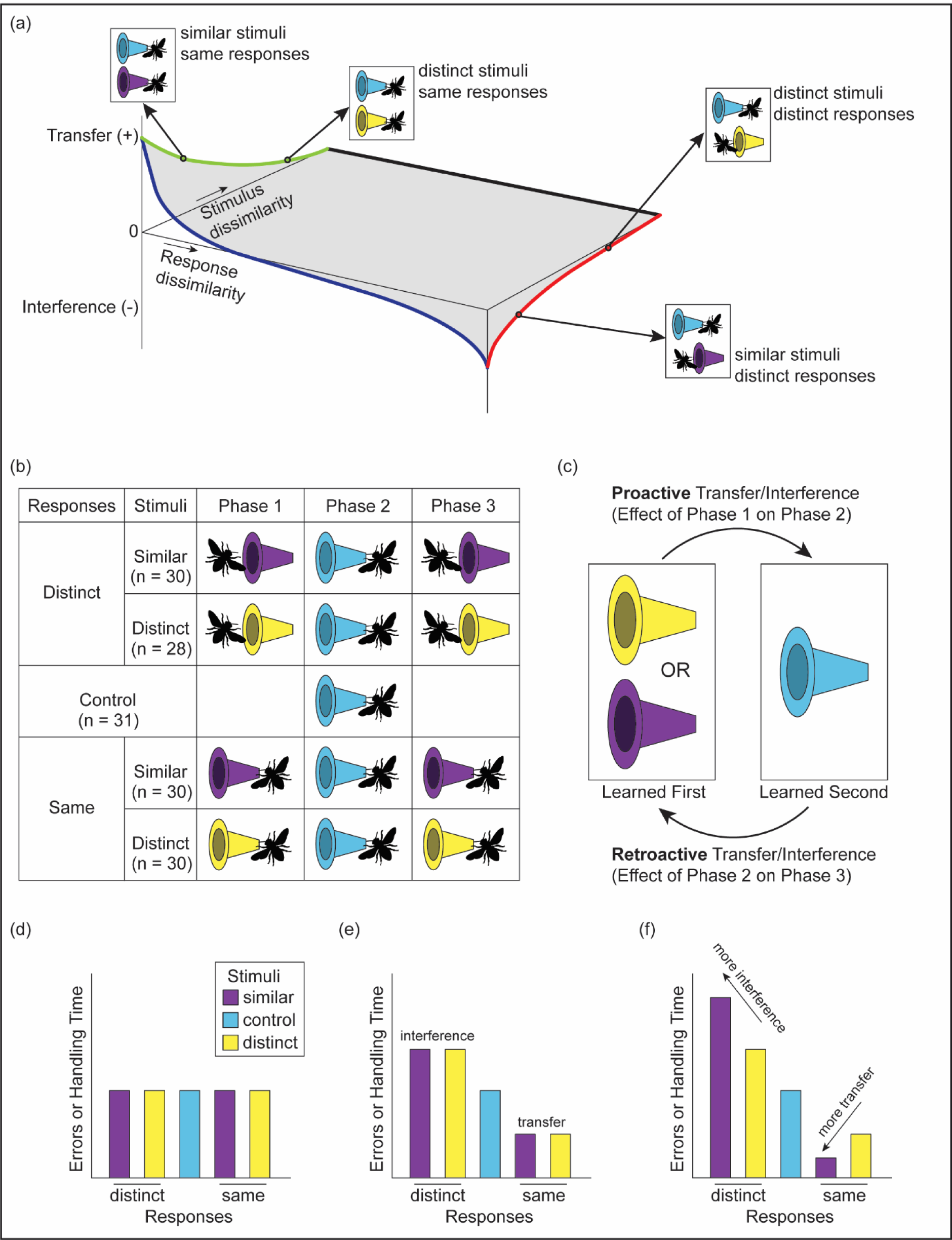
(a) The relationship between stimulus dissimilarity, response dissimilarity, and expression of transfer and interference, as based on Osgood’s transfer surface model. Specific examples of stimulus – response pairs are given at each point for clearer understanding. (b) Summary of five treatments used in the experiment. A bee at the front of a flower represents legitimate visitation, and a bee at the back of a flower represents nectar robbing. (c) Schematic view of retroactive and proactive effects. (d) Predicted outcomes for the treatment groups when neither response similarity nor stimulus similarity affects performance, (e) when only response similarity affects performance, and (f) when both response similarity and stimulus similarity interact to affect performance, based on Osgood’s transfer surface model. More errors or higher values of handling time represent lower performance.

Considering the importance of stimulus similarity and response similarity in understanding how sequentially-formed memories interact, no study has investigated the interaction of stimulus and response similarity in animals. Studies to date have manipulated either stimulus similarity or response similarity, but not both. For example, a greater similarity in responses was found to result in greater transfer in a study of crabs (Hughes & O’Brien, 2001); however, stimulus similarity was not evaluated as a factor. In studies of birds and rats where stimuli in learned tasks were similar but responses were distinct, there was significant interference (Lewis & Kamil, 2006; Roberts, 1981). There is also a substantial body of literature on the transfer of knowledge to novel stimuli, although it is often not explicitly stated as transfer. In such cases, when novel stimuli resemble old ones, there tends to be greater generalization (Ghirlanda & Enquist, 2003). For example, animals express previously learned antipredator responses to a novel predator if it resembles the predator they encountered before, that is, if stimuli associated with different predators were similar (Brown et al., 2011; Ferrari et al., 2007; Ferrari & Chivers, 2009; Griffin et al., 2001). However, these evaluations of stimulus similarity did not consider the role of response similarity.

In this study, we explored Osgood’s model with respect to nectar foraging behaviour in the common eastern bumble bee, *Bombus impatiens*. Bumble bees are ideal organisms for investigating how stimulus similarity and response similarity influence transfer and interference in learning and memory. Bumble bees use a variety of flowers in the wild and associate traits of those flowers (e.g., colour, scent, pattern or shape) with rewards such as nectar (Clarke et al., 2013; Dyer et al., 2006; Giurfa et al., 1996; Gumbert, 2000; Whitney et al., 2009; Wright & Schiestl, 2009). Accessing nectar from flowers require different motor routines depending on flower morphology (Krishna & Keasar, 2021; Laverty, 1994b). Individual bumble bees can learn how to handle more than two types of flowers that require different handling techniques (Chittka & Thomson, 1997; Gegear & Laverty, 1998; Ishii & Kadoya, 2016; Krishna & Keasar, 2019; Woodward & Laverty, 1992). In short, both stimuli (e.g., colour, scent, pattern or shape) and responses (e.g., motor routine, flower choice) vary among flowers used by these bees.

Transfer and interference in bees and butterflies have been studied in multiple contexts. These include serial reversal learning (Mota & Giurfa, 2010; Strang & Sherry, 2014), navigation (Cheng, 2005; Cheng & Wignall, 2006), learning the cue that indicates nectar (Dukas, 1995), nest (Colborn et al., 1999; Fauria et al., 2002; Worden et al., 2005), or oviposition site (Weiss & Papaj, 2003). Much research on transfer and interference has focused on learning how to handle flowers to get nectar rewards (Chittka & Thomson, 1997; Gegear & Laverty, 1998; Goulson et al., 1997; Krishna & Keasar, 2021; Laverty, 1994a, 1994b; Lewis, 1986; Lichtenberg et al., 2020; Raine & Chittka, 2007; Woodward & Laverty, 1992). Interference between different learned flower handling motor routines has been proposed to explain why pollinators often show flower constancy, a behaviour where pollinators consistently forage on a single flower species even though alternative resource is available (Chittka et al., 1999; Grüter & Ratnieks, 2011). Importantly, no study to our knowledge has examined the interaction between floral stimulus similarity and similarity in flower handling routines.

In our study of flower foraging, we evaluated two specific assertions of Osgood’s model: 1) response similarity determines whether a response will be transferred or will interfere with other responses and 2) stimulus similarity determines the magnitude of the transfer or interference effect through generalization. To evaluate these assertions, we required bumble bees to learn two stimulus – response associations sequentially and manipulated stimulus similarity (in terms of flower colour) and response similarity (in terms of flower handling motor routine) within each association pair (summarized in Figure 1b). Our evaluation of response similarity focused on two types of flower handling techniques: legitimate visitation and nectar robbing. Legitimate visitation involves bees probing into or entering through the flower opening to access the nectar reward. Nectar robbing occurs when bees access nectar through a hole at the base of the flower that they or other individuals have created (Inouye, 1983; Irwin et al., 2010). We measured proactive effects (from the first to the second learned association) and retroactive effects (from the second to the first learned association) using flower handling time and the number of failed flower visits (Figure 1c). Additionally, we recorded where bees landed on a flower during failed visits to determine if errors resulted from incorrectly applying a previously learned response.

We tested three specific alternative predictions: 1) If neither response similarity nor stimulus similarity affects performance, experienced bees will show the same flower handling time/failed flower visits as control bees with no prior experience (Figure 1d); 2) If only response similarity determines whether the learned association is transferred and interfered with, and stimulus similarity has no effect, bees that experienced distinct responses will show interference, while bees that experienced same responses will show transfer, regardless of flower colour similarity (Figure 1e). 3) If bee behaviour fits the Osgood’s model, response similarity and stimulus similarity will interact. Thus, if motor routines for novel and previously-learned associations are distinct, bees will show greater interference when flower colour in sequential association is more similar. This interference will be due to novel stimuli incorrectly eliciting the previously learned response. In contrast, when the motor routine required for the novel association is the same as the previously-learned association, bees will show greater transfer when flower colour is more similar (Figure 1f).

## Methods

### Bumble bee colony maintenance

Bumble bee colonies were purchased from Koppert Biological Systems (Howell, MI, USA). A total of 149 worker bees from three colonies were used for the experiment. Each colony (nest box dimension: 25 × 23 × 12 cm) was connected to two separate chambers constructed from plywood via Tygon tubing. The dimensions of these chambers were 82 × 60 × 60 cm (large) and 38 × 60 × 40 cm (small), respectively. A 20% (v/v) sucrose solution was provided ad libitum in both chambers in 148 ml (40 dram) vials (BioQuip Products Inc., USA), with a lid. A hole was made in the lid, through which a cotton wick (15 cm braided cotton rolls; Richmond Dental Inc.) was inserted to continuously draw the sugar solution to the top of the vial. To encourage bees to fly to the feeder, rather than walk, feeders were suspended in mid-air using wires. Only the large chamber was used for the behavioural assay. During the assay, access to the large chamber was restricted to a test bee only, while other bees were still allowed to forage on the small chamber. Ground honeybee corbicular pollen (Koppert Biological Systems, MI, USA) was directly supplied to the colony daily. Both chambers were painted grey (Glidden, PPG Industries Inc., PA, USA) on their walls and had a thin layer of beige sand on the floor. A transparent window was placed on one side of each chamber to facilitate observation. The chambers’ ceilings were made of 3 mm acrylic plastic, with 40 W 4400 lumen LED light panels (61 × 61 cm LED Panel; 5000 K Cool White, James Industry) positioned overhead. The lighting was regulated by a timer, maintaining a 12:12 hour light:dark cycle.

### Feeder and artificial flowers design

We used an array consisting of 4 vertical rods, each spaced 10 cm apart horizontally. On each rod, 4 feeders were attached, each spaced 10 cm apart vertically. This 4 × 4 vertical array design (total 16 feeders) allowed bees to approach a feeder from the back as well as the front. A feeder was composed of two syringes connected by a tube filled with water, a small wire loop connected to one of the syringes, and a sucrose solution reservoir (Figure 2a). When loaded, the small wire loop holds 1.95 ± 0.17 µL (mean ± SD) of 50% (v/v) sucrose solution by surface tension. By pulling and pushing one syringe, hydraulic force caused the syringe on the other side to move up and down. The wire loop would submerge into the 50% sucrose solution reservoir as the syringe moved down and replenished the sucrose solution. The syringes used to operate the feeder were located outside of the arena. In this way, the feeder design allowed manual replenishment of the feeder without disturbing a focal bee. The entire array and feeders were painted in the same grey colour as the chamber walls. During the experiment, artificial flowers were mounted on the feeders (Figure 2b).

**Figure 2.**
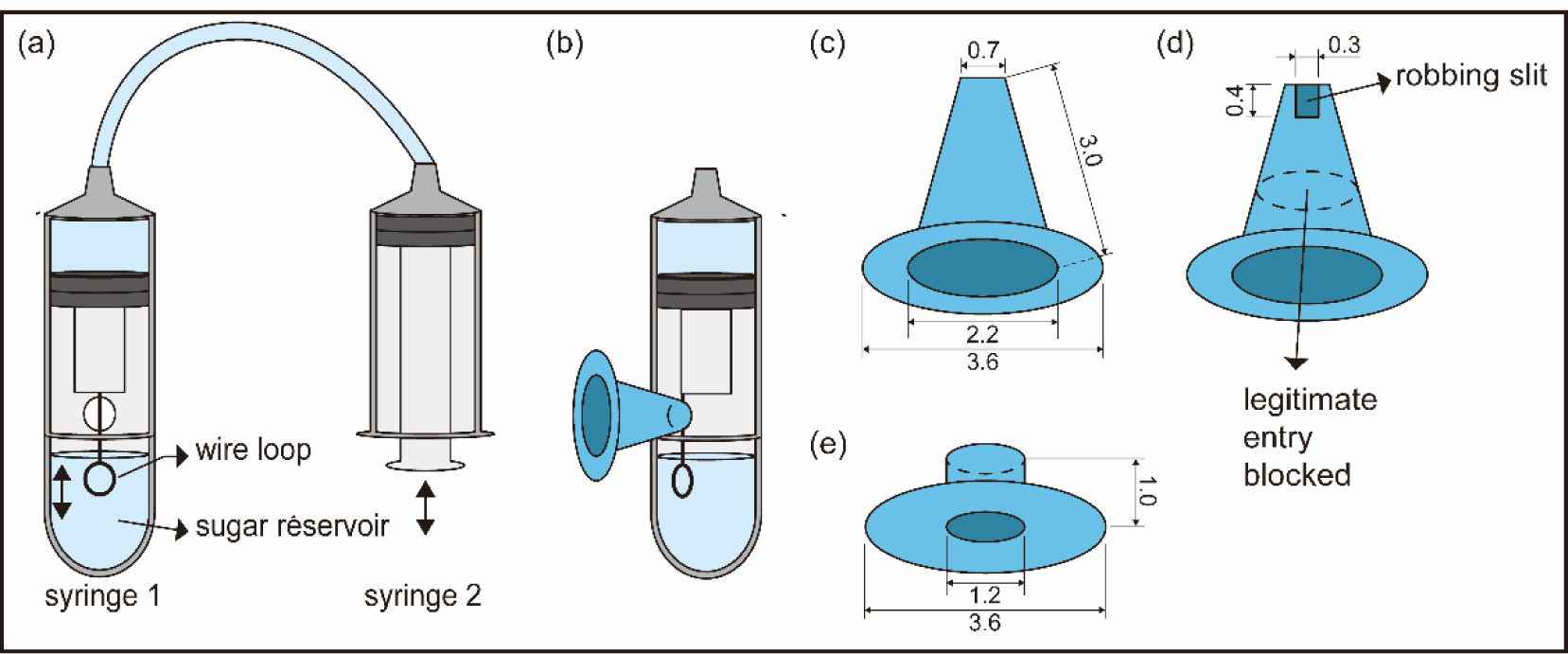
Diagram illustrating the design of the feeder and artificial flowers. (a) Hydraulic force operated feeder. Manipulating syringe 2 controls the vertical movement of the wire loop connected to syringe 1. (b) Artificial flower mounted on a feeder. (c) Flower accessible for legitimate visits. (d) Flower accessible for nectar robbing. (e) Pre-training flower.

The artificial flowers used in the experiment consisted of two parts: a circular petal section and a truncated cone-shaped section, which allows a bee to enter inside of a flower (Figure 2c and d). Flowers for the pre-training phase also consisted of two parts: a circular petal section and a short cylinder-shaped section (Figure 2e). To create flowers in either blue, purple, or yellow (Figure 3a), thick coloured paper (Canson Colorline paper 150 gsm, Canson, France) was used. The combination of yellow and blue represent distinct stimuli, while purple and blue represent similar stimuli. Each coloured flower could be handled either by legitimate visit or nectar robbing, but not both. Flowers that could be handled by legitimate visit did not have robbing holes, allowing only legitimate visits from its entrance (Figure 2c). Flowers that could be handled by nectar robbing possessed a 3 × 4 mm robbing hole on the basal part of the cone, but access via the legitimate entrance of the flower was blocked by a transparent acrylic sheet (Figure 2d). The combination of legitimate visit and nectar robbing represent distinct responses, and constant robbing represents same responses.

**Figure 3.**
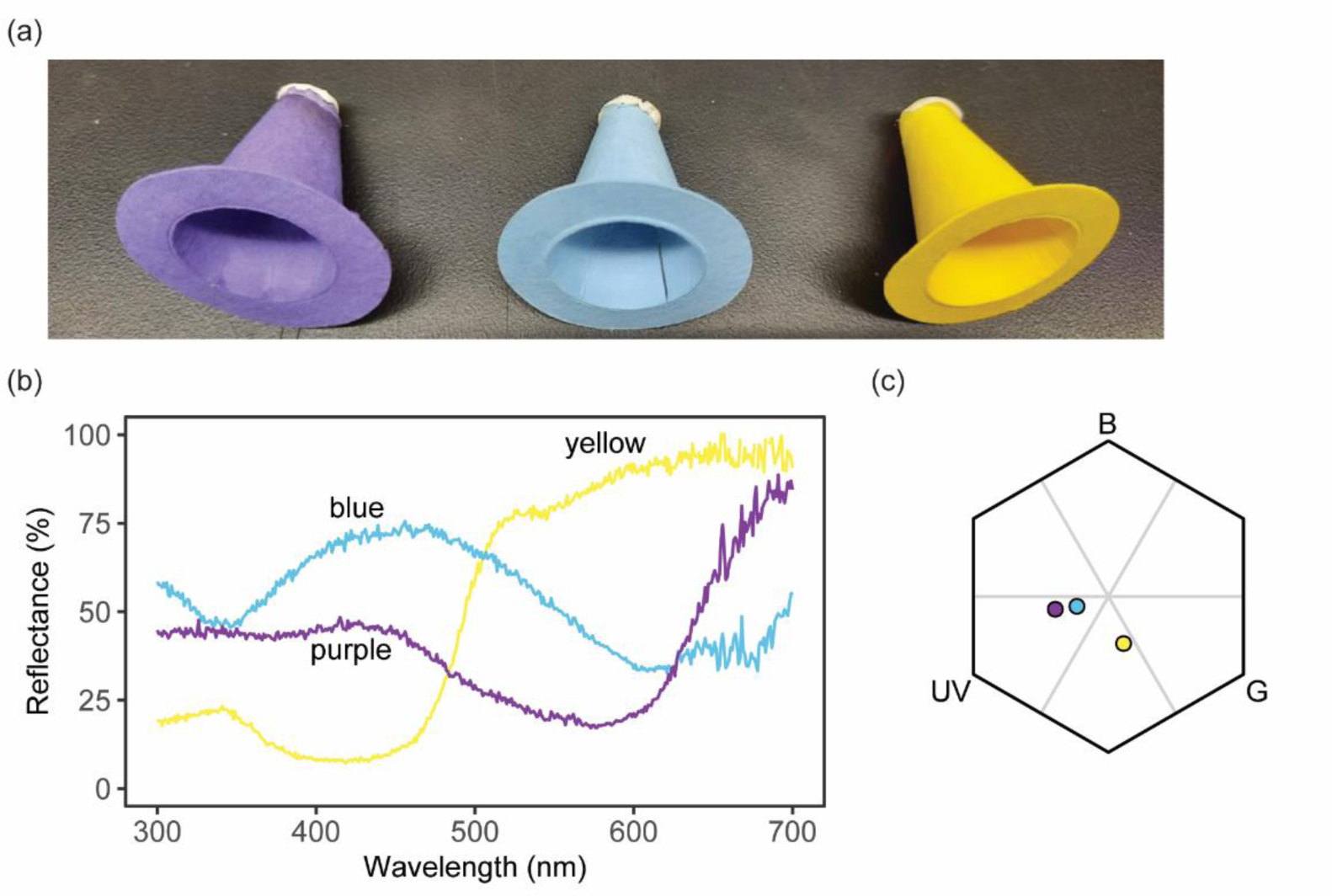
Colour differences of artificial flowers used in the experiment. (a) The three colours of artificial flowers used (b) The spectra of flower colours (c) The flower colours placed in bumble bee colour space.

The reflectance spectrum of each flower colour was measured using an UV-VIS spectrophotometer (Ocean Optics USB2000, USA) with deuterium-tungsten halogen light source (Ocean Optics DH2000, USA), and an integration time of 3 ms (Figure 3b). To ensure that the colours were distinguishable by bees, we employed the hexagon colour model for bee vision (Chittka, 1992) based on the spectral sensitivities of *B. impatiens* (Skorupski & Chittka, 2010). We plotted the locus of each flower colour within the hexagon colour space (Figure 3c). Reflectance spectra were converted into colour loci by calculating the excitation of three types of colour receptors. For the calculation, we used the chamber’s illumination and grey (the wall of chamber) as the background. The Euclidean distance between purple and blue was determined to be 0.14 units, while the distance between yellow and blue measured 0.38 units. Bumble bees have been reported to be capable of discerning colours more than 0.045 units apart from one another (Dyer & Chittka, 2004). Thus, all colours used could presumably be discriminated against by the bees, but yellow was nevertheless more different from blue than purple was from blue.

### Experiment protocol

During the pre-training phase, yellow and purple pre-training flowers were presented to multiple bees for a duration of 1-2 hours. The purpose was to motivate the bees to visit cone-shaped artificial flowers in the subsequent testing phase. The pre-training flowers contained 50% sucrose solution, which was replenished using eye droppers. Bees that visited both the yellow and purple pre-training flowers were marked with colour paint markers (POSCA, Mitsubishi pencil CO.) using unique colour combinations for individual identification.

Pre-trained bees went through three subsequent learning phases. Only one bee was allowed in the experiment chamber (the large chamber) at a time, where it foraged freely on a 4 × 4 vertical array of flowers. Each bee was given an array consisting of a single type of flower, with varying stimulus and response for each phase (summarized in Figure 1b):

<u>Phase 1:</u> single type of flower array.
(stimulus: purple or yellow, response: legitimate or robbing visit)
<u>Phase 2:</u> novel type of flower array.
(stimulus: blue, response: robbing visit)
<u>Phase 3:</u> reintroduction of the same type of flower array as in phase 1.

This experimental design allowed us to investigate both the proactive and retroactive effects of sequentially learned stimulus – response associations. The performance on novel association learned in phase 2 shows the proactive effect of the association previously learned in phase 1. The performance on the association reintroduced in phase 3 shows the retroactive effect of novel association learned in phase 2 (Figure 1c).

Each bee was randomly assigned to one of five treatment groups (four experimental groups and a control group). All groups were given blue robbing-accessible flowers as novel flowers in phase 2, but they differed in flower type received in phases 1 and 3, which varied in colour (purple or yellow), and in the motor routine required to access nectar (legitimate visit or nectar robbing). Thus, stimulus similarity and response similarity associated with those stimuli varied across treatments, creating a 2 × 2 factorial design of “similar stimuli” vs. “distinct stimuli” and “same responses” vs. “distinct responses” (Figure 1b). In the “similar stimuli” treatments, bees experienced purple flowers in phase 1, blue flowers in phase 2, and purple flowers again in phase 3. In the “distinct stimuli” treatments, bees experienced yellow flowers in phase 1, blue flowers in phase 2, and yellow flowers again in phase 3. In the “distinct responses” treatments, responses associated with stimuli were different across the phases. In this case, yellow or purple flowers were only accessible through legitimate visits. Conversely, blue flowers were only accessible through robbing. Thus, bees were required to visit legitimately in phase 1, were required to rob in phase 2, and then required to visit legitimately again in phase 3. In “same responses” treatments, responses associated with stimuli were the same across the phases. Specifically, access to sucrose required robbing throughout all three phases. The control group of bees experienced phase 2 (blue flowers with a robbing hole) only without going through phases 1 and 3.

Throughout these phases, once bees filled their crop, they were allowed to return to their colony to empty it. Bees that successfully visited flowers more than 30 times in a given phase proceeded to the next phase the next time they left the colony. A successful visit was defined as a bee spending more than one second after inserting proboscis into the robbing hole of a flower accessible for robbing or after entering the legitimate entrance of a flower accessible for legitimate visit. Most bees achieved 30 successful visits within a single foraging bout, but some required two bouts to meet the criteria. All bees finished three phases within a day. Behavioural observations were recorded using a GoPro camera (GoPro Hero 5, GoPro Inc., San Mateo, CA, USA) at 30 frames per second.

During phases 2 and 3, the flower handling time (time from landing on a flower to leaving) and the number of failed visits (instances where a bee landed on a flower but left without obtaining a sucrose reward) until 30 successful visits were measured as proxies of learning performance by using BORIS software (Friard & Gamba, 2016).

### Statistical Analyses

To ensure consistency among all bees that completed the assay, only measurements of the first 30 successful visits were analysed, as the total number of successful visits varied between bees (Number of successful visits; Phase 1: mean = 49.7, SD = ± 11.2; Phase 2: mean = 52.0, SD = ± 11.7; Phase 3: mean = 47.1, SD = ± 10.0) and performance could be a function of the number of successful visits. Flower handling time was averaged for every consecutive block of 10 visits, resulting in three blocks for each phase.

To investigate the proactive and retroactive effect of learned stimulus – response associations on bees’ flower handling performance, we analysed the effect of flower treatments on two key variables in phase 2 and 3: averaged flower handling time and the number of failed visits to flowers. For phase 2, we fitted a generalized linear mixed model with the number of failed visits to flowers as a dependent variable, assuming a negative binomial distribution to account for overdispersion. The type of previous flower bees experienced was a fixed effect, and colony identity was a random effect. Then, we conducted a linear mixed model with the averaged flower handling time as a dependent variable, and the order of blocks of visits, type of previous flowers bees experienced, and their interaction term as fixed effects. Averaged flower handling time was log-transformed for further analysis to improve the normality of the residuals. Colony identity and bee identity were included in the model as random effects. Same models were fitted for phase 3, but the interaction term in the linear mixed model for average flower handling time was dropped after it was checked as non-significant.

To evaluate whether failed visits were resulted from generalization of previously learned and novel colours, and thus applying the previously rewarding handling technique to novel flowers, failed visits were further categorized based on whether the bee landed on the front part of the circular petal section (failed visits on the front); or the back part of the circular petal section or the truncated cone shape section (failed visits on the back). We fitted the separate GLMMs for those two types of failed visits. The same fixed effect and random effect as the model for total failed visits are used for GLMM with negative binomial distribution.

All statistical analyses were done by R version 4.3.1 (R Core Team, 2023). We ran GLMMs and LMMs using the lme4 package (Bates et al., 2015). To calculate significance of fixed effects in models, we compared the models with and without the focal fixed effects using the parametric bootstrap with 1000 replicates by the PBmodcomp function in pbkrtest package (Halekoh & Højsgaard, 2014). Tukey post-hoc pairwise comparisons were performed with the emmeans function in emmeans package (Lenth, 2022).

### Ethical Note

We followed all the legal requirements of the U.S. and all institutional guidelines to maintain the welfare of bumble bees during this research. The colonies used in this research were kept under suitable temperature conditions and provided with an ample supply of food. Bees were humanely euthanized by freezing at the end of the experiment.

## Results

### Proactive effects: Do stimulus and response similarities between previous task and novel task affect performance on the novel task?

In phase 2, bumble bees’ performance on novel flowers (i.e., blue flowers that required a robbing technique) varied according to similarities in flower colours and motor routines between previously experienced flowers and novel ones (Figure 4a). When bees were forced to switch their handling technique, from purple to blue flowers, they made more failed visits than naïve bees (Tukey post-hoc comparison: log-estimate ±SE = −1.14±0.14, *P* < 0.0001), demonstrating a proactive interference. Switching handling technique, from yellow to blue flowers, did not show interference or transfer compared to naïve bees (Tukey post-hoc comparison: log-estimate ±SE = −0.19±0.15, *P* = 0.71). Conversely, bees allowed to use the same handling technique made fewer failed visits than naïve bees, demonstrating a proactive transfer (Tukey post-hoc comparison: control vs. “distinct stimuli – same responses”, log-estimate ±SE = 0.44±0.15, *P* = 0.034; control vs. “similar stimuli – same responses”, log-estimate ±SE = 0.98±0.17, *P* < 0.0001). Transfer effects were greater when the colour of novel flower was similar to that of the flower they had experienced earlier in phase 1.

**Figure 4.**
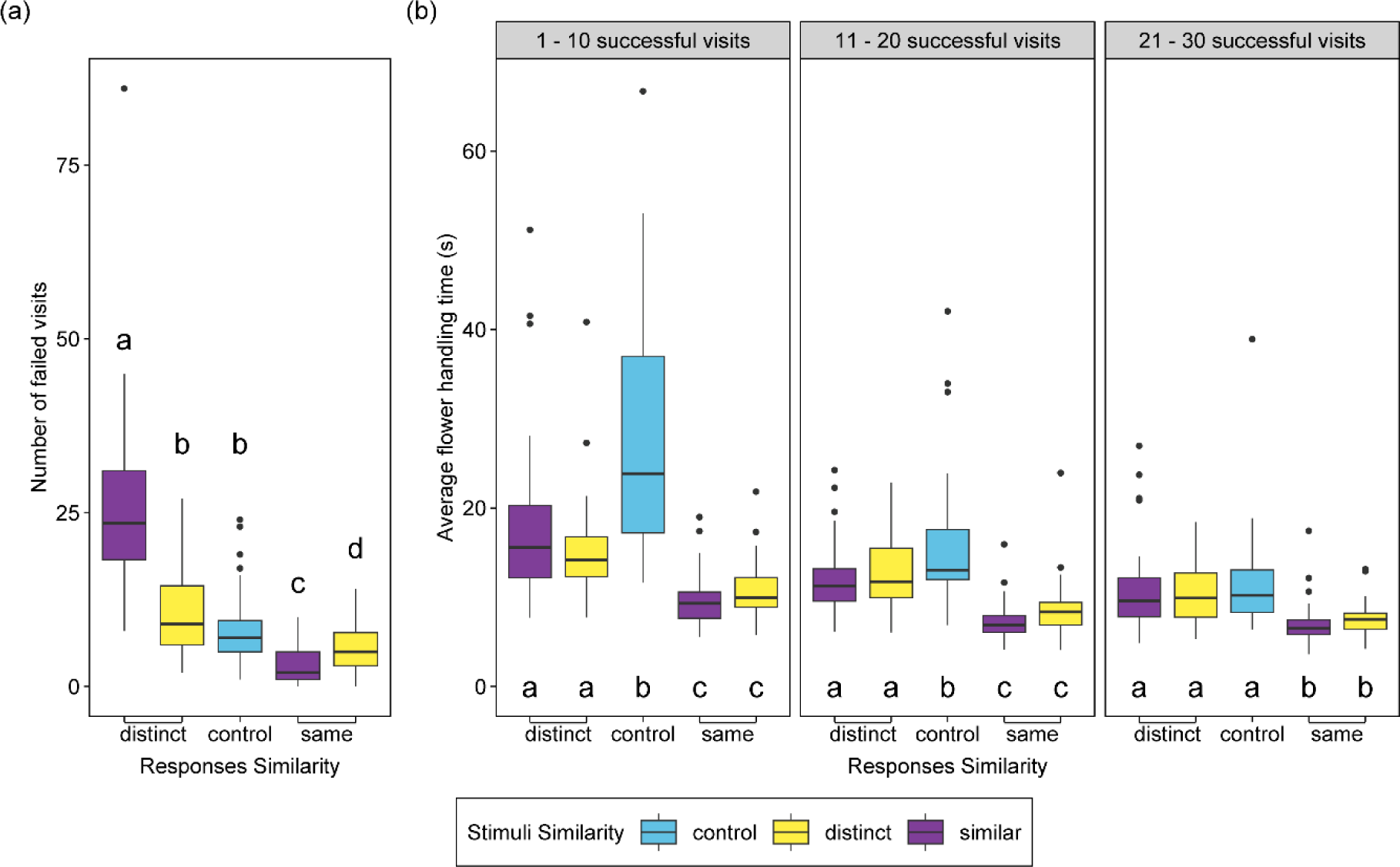
Bumble bees’ performance on novel flowers during Phase 2. (a) Number of failed visits to flowers, and (b) Flower handling time averaged per 10 successful visits across the blocks of visits. Box plots represent medians, 25th and 75th percentiles. Whiskers extend to the farthest data points within 1.5 times the interquartile ranges. Dots represent data points outside 1.5 interquartile ranges. Letters indicate groups within each graph panel that are significantly different at *P* < 0.05 from Tukey post hoc test.

Flower handling time was significantly affected by the order of consecutive blocks of 10 visits and type of previous flowers bees experienced (LMM with log transformation: *χ^2^_4_* = 48.83, *P* < 0.0001). Bees became faster in handling flowers as they completed more successful visits to flowers (LMM with log transformation: estimate ±SE = –0.41±0.03, *P* < 0.0001). Bees that had learned the legitimate technique experienced longer flower handling times when they switched to nectar robbing compared to those that had previously learned nectar robbing technique, yet they were faster than naïve bees (Figure 4b). Therefore, regardless of their previously learned technique, all experienced bees showed a proactive transfer. However, stimulus similarity did not affect the degree of this transfer effect. The differences in flower handling time among “distinct responses” and “control” treatments disappeared in the third block of 10 successful visits (21-30 visits) to flowers (Figure 4b).

### Retroactive effects: Do stimulus and response similarities between previous task and novel task affect the performance on the previously learned task?

In phase 3, we investigated whether bees exhibited changes in performance when they switched back to the original flower type that they experienced in phase 1. Unlike the proactive effects, retroactive effects in the third phase cannot be interpreted as transfer or interference due to the limitation of our experimental design. To rigorously investigate whether the newly learned association enhances or interferes with the previously learned association, performance of the experimental group would have to be compared with a control group that exclusively foraged on a single flower type, which was absent in our design (Chittka & Thomson, 1997; Woodward & Laverty, 1992).

The number of failed visits was significantly different among treatments (Figure 5a; GLMM with negative binomial distribution: *χ^2^* = 217.36, *P* = 0.001). Bees required to switch back from robbing to legitimate visits made more failed visits to purple flowers than yellow flowers, indicating the effect of stimulus similarity (Tukey post-hoc comparison: log-estimate ±SE = 1.36±0.18, *P* < 0.0001). However, bees that only nectar robbed in each phase did not show a difference in their failed visits based on stimulus similarity (Tukey post-hoc comparison: “similar stimuli – same responses” vs. “distinct stimuli – same responses”, log-estimate ±SE = 0.19±0.23, *P* = 0.85).

**Figure 5.**
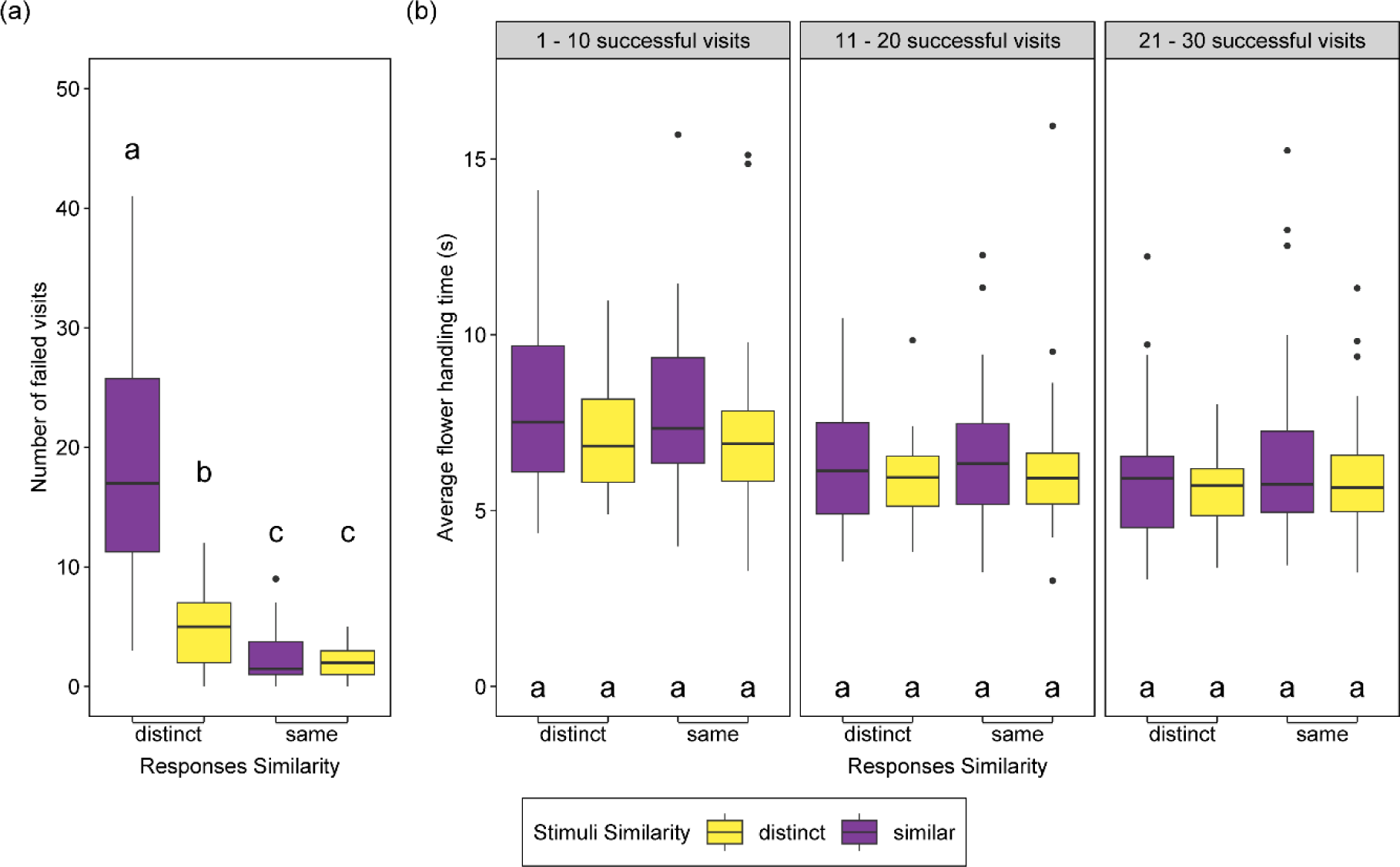
Bumble bees’ performance on flowers during Phase 3. (a) Number of failed visits to flowers, and (b) Flower handling time averaged per 10 successful visits across the blocks of visits. Box plots represent medians, 25th and 75th percentiles. Whiskers extend to the farthest data points within 1.5 times the interquartile ranges. Dots represent data points outside 1.5 interquartile ranges. Letters indicate groups within each graph panel that are significantly different at *P* < 0.05 from Tukey post hoc test.

Flower handling time was not affected by treatment type, and only the order of consecutive blocks of 10 visits had a significant effect (Figure 5b; LMM with log transformation: the order of blocks, *χ^2^_1_* = 138.77, *P* = 0.001; treatment, *χ^2^_3_* = 1.62, *P* = 0.66). Regardless of the type of flowers, bees become faster in handling flowers over time (Figure 5b; LMM: estimate ±SE = –0.11±0.008, *P* < 0.0001).

### Failed visits according to the landing positions on flowers

Further categorizing failed visits based on the landing positions on flowers by bees revealed that differences in failed visits among treatments were due mainly to the type of motor routines that were rewarding in the previous phase, suggesting generalization of previous-acquired knowledge into a novel context. In phase 2, the number of failed visits from the front side of the flower showed a similar pattern to the overall failed visits (GLMM with negative binomial distribution: *χ^2^* = 252.75, *P* = 0.001). Bees with previous experience of legitimate visits had a higher frequency of failed visits from the front side than bees with previous experience of robbing, and this effect was greater in the “similar stimuli” treatment (Figure 6a). Although there was a significant treatment effect for failed visits from the back of a flower (GLMM with negative binomial distribution: *χ^2^* = 14.88, *P* = 0.003), changes in stimulus similarity did not affect the number of failed visits (Figure 6b).

**Figure 6.**
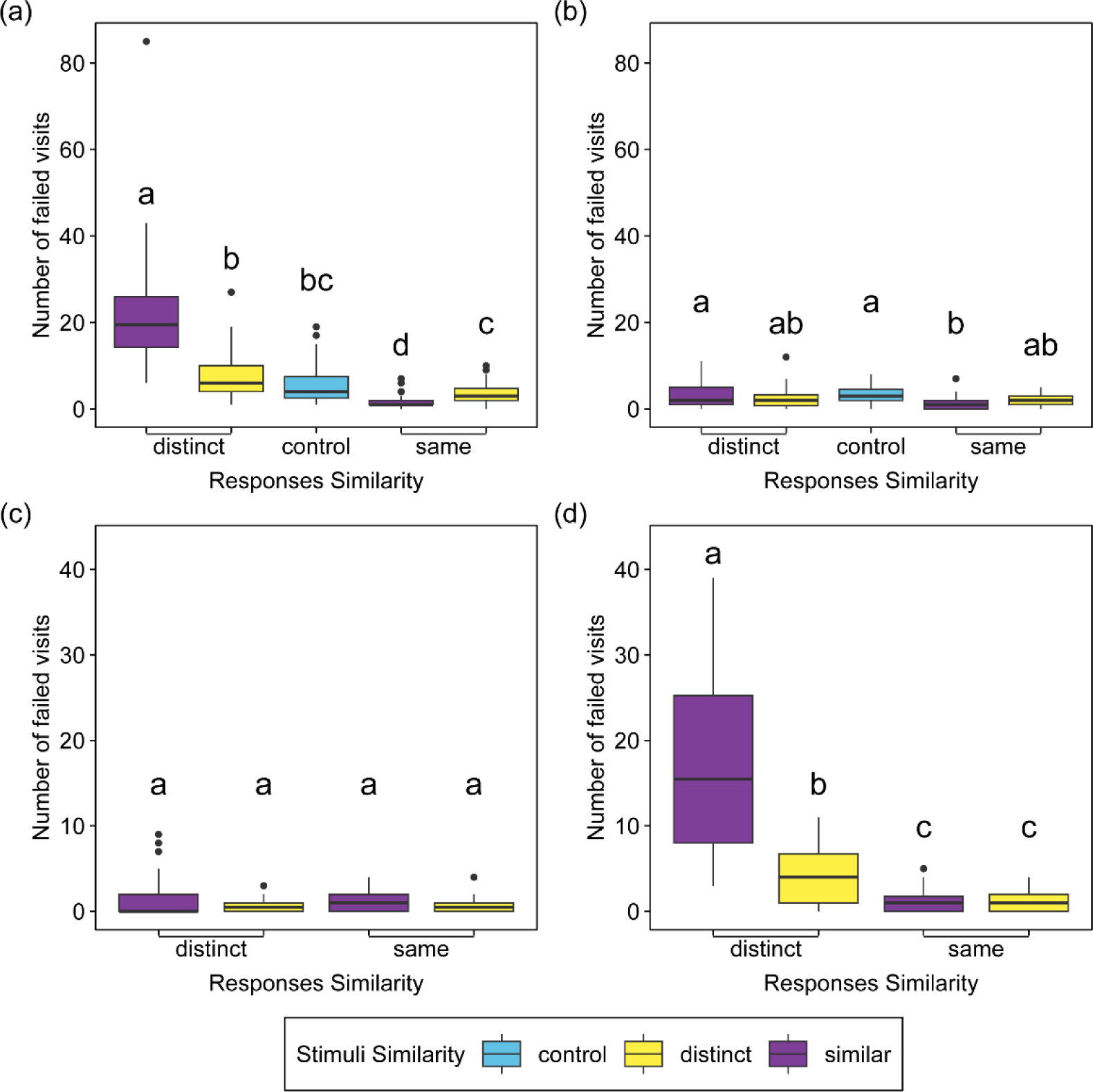
Comparison of failed visits to flowers in phases 2 and 3 from different landing positions. (a) Failed visits when bees landed on the front side, and (b) on the back side of a flower in phase 2. (c) Failed visits when bees landed on the front side, and (d) on the back side of a flower in phase 3. Box plots represent medians, 25th and 75th percentiles. Whiskers extend to the farthest data points within 1.5 times the interquartile ranges. Dots represent data points outside 1.5 interquartile ranges. Letters indicate groups within each graph panel that are significantly different at *P* < 0.05 from Tukey post hoc test.

In contrast, during phase 3, bees did not differ in the frequency of failed visits from the front side of a flower regardless of the type of flowers they handled (GLMM with negative binomial distribution: *χ^2^* = 5.82, *P* = 0.14; Figure 6c). Instead, failed visits from the back side of a flower showed a consistent pattern with the overall failed visits (GLMM with negative binomial distribution: *χ^2^_3_* = 255.72, *P* = 0.001; Figure 6d).

## Discussion

Our study manipulated both the degree of similarity in flower stimuli and similarity in flower handling responses experienced by bumble bees during foraging, aiming to elucidate their effects on transfer and interference of sequentially learned stimulus – response associations. We found a significant interaction between stimulus and response similarities on failed visits to flowers, which was consistent with Osgood’s model; in contrast, we did not find the interaction in terms of flower handling time. To our knowledge, this finding represents the first comprehensive examination of Osgood’s transfer plane model in nonhuman animals. We are unaware of any study of nonhuman animals showing an interaction between stimulus similarity and response similarity in the expression of transfer and interference.

### Proactive interference

In our study, proactive interference occurred when bees incorrectly applied the motor routine that had been rewarding in the first phase to a flower requiring the alternative motor pattern in the second phase. Notably, interference was observed only in failed visits for bees assigned to the similar stimuli – distinct responses treatment. This is the condition predicted in Osgood’s model to result in greater interference; however, even in this treatment, flower handling time was unexpectedly shorter than that of naïve bees. Proactive interference has been observed in other studies of insects, including bees (Chittka & Thomson, 1997; Dukas, 1995; Laverty, 1994b). Moreover, the lack of interference in the other treatments is consistent with the existing literature, which has generally shown little or no interference when bees switch between different flowers (Chittka & Thomson, 1997; Krishna & Keasar, 2021; Laverty, 1994a; Lichtenberg et al., 2020; Raine & Chittka, 2007; Woodward & Laverty, 1992). In those previous studies, the effect of similarity on interference was not evaluated. Our study underscores the importance of considering both the stimulus similarity and the response similarity when assessing interference, as well as the choice of metrics for measuring foraging performance. For example, Dukas (1995) found that prior experience in discriminating a rewarding colour from an unrewarding one could result either in interference or in transfer to the later task, depending on the specific colour pair being discriminated. Osgood’s model would predict that this dependence was due to instances where the rewarding colour was similar to either the rewarding or the unrewarding colour in the subsequent phase.

### Proactive transfer

Transfer effects have also been observed in bees (Chittka & Thomson, 1997; Dukas, 1995; Laverty, 1994b). For example, bees that became experienced in handling simple structured flowers showed decreased handling time of other simple structured flowers (Laverty, 1994b). Similar transfer effects have also been documented in other animals, where prior experience in solving a functionally identical, but perceptually different problem, aided in selecting the right tool to solve the subsequent problem (Bobrowicz et al., 2020, 2021). The degree of transfer tends to be greater when responses are similar or when stimuli are similar (Cheng, 2005; Colborn et al., 1999; Fauria et al., 2002; Lewis et al., 2013; Roberts, 1981). Our findings regarding failed visits align with this observed pattern and with Osgood’s framework, as we observed that transfer increased more in similar stimuli – identical responses treatment than distinct stimuli – identical responses treatment. However, this pattern was not the same for flower handling time, as bees displayed transfer irrespective of the similarity in responses and stimuli.

### Different results between flower handling time and failed visits to flowers

The inconsistency between failed visits and flower handling time could be attributed to the similarities in the motor routines required for handling flowers in our experiment. We initially assumed that legitimate visitation and nectar robbing involve different flower handling techniques. However, our results suggest that the degree of similarity between these two techniques changes as the motor routines play out. For example, once bees landed on the correct side of a flower, where the legitimate entrance or nectar robbing slit were located, they only needed to insert their proboscis into a hole to obtain sugar reward irrespective of the flower handling technique they used. Thus, bees employing both legitimate and nectar robbing techniques share components of motor routines such as proboscis insertion, with neither technique interfering with the other in that regard. As a result, bees might experience interference when deciding to land on which side of a flower, but experience transfer later in terms of time spent collecting nectar on a flower.

This interpretation aligns with observations of Chittka and Thomson (1997), who found that bees made more errors when switching between different motor tasks, even though the handling times decreased. As in our study, bees were required to select the rewarding direction (left or right), and once the decision was made, the subsequent flower handling process was the same in both tasks. Furthermore, transfer of required skills has been shown to vary over the course of behaviour in studies of primates and birds (Chittka, 1998; Chittka & Thomson, 1997; Keasar et al., 1996; Laverty, 1994b; Mirwan et al., 2015). When they were required to solve the novel task using a tool, transfer effects occurred during the early stages of executing a new behavioural sequence, however, the overall performance was not affected.

Alternatively, contextual information provided after landing on a flower may have reduced interference in flower handling time. In our study, basic similarities in the flower structure itself may have helped bees to minimize confusion in executing the appropriate motor routine. Contextual cues are known to reduce interference effects (Cheng, 2005; Colborn et al., 1999; Fauria et al., 2002; Lewis et al., 2013; Roberts, 1981), unless the contextual cues are not sufficiently learned (Fauria et al., 2002).

### Retroactive effects

Retroactive effects on the previously learned stimulus – response association in our study were not as pronounced as proactive effects. In the third phase, where retroactive effects were assessed, differences in flower handling times disappeared, with all bees showing short handling times. Bees still made more failed visits to purple flowers than yellow flowers when responses were distinct, but they did not show a difference in failed attempts based on flower colour when responses were identical. One plausible explanation for this pattern is that the learning performance of nectar robbing reached an asymptote by the third phase for both colours, as shown in other studies of bumble bees handling flowers (Chittka, 1998; Chittka & Thomson, 1997; Keasar et al., 1996; Laverty, 1994b; Mirwan et al., 2015). By this point, bees that only nectar robbed in each phase might have gained sufficient experience resulting in a small number of failed attempts, irrespective of flower colour.

### Dealing with multiple flower species

When both nectar robbing and legitimate visitation are possible on a single flower species, insect pollinators tend to select the tactic that maximizes energy efficiency (Krishna & Keasar, 2019; Laverty, 1994b). However, most studies of nectar robbing have focused on a single plant species. The decision-making process regarding nectar robbing and legitimate visitation on multiple flower species has not yet been investigated, even though individual bees may encounter various flower species over their lifetime and even during a single foraging bout (Ishii & Kadoya, 2016). Tactic constancy, defined as the tendency of bees to employ the same flower handling tactic even when alternative tactics are available, has been proposed as a strategy to minimize interference that may arise from switching tactics (Bronstein et al., 2017). Moreover, it has been suggested that the similarity between motor routines involved in flower handling influences bees’ switching patterns (Ishii & Kadoya, 2016). We propose that nectar robbing on various flower types is more transferable than legitimate visitation on the same flower types. Legitimate visitation on one flower type may require a quite different handling routine than the one used earlier, owing to the diversity of complex morphologies that flowers have (Krishna & Keasar, 2019; Laverty, 1994b). In fact, the diversity of floral morphology has been suggested to have been favored by selection to ensure that bees remain faithful longer to a given plant species (Schiestl & Johnson, 2013; Wei et al., 2021). In contrast, nectar robbing techniques seem more similar across different flowers, as the presence of the nectar robbing hole bypasses the complex morphology of the floral entrance.

In addition to nectar foraging, bees also collect pollen to raise their young, thus expanding the possible number of motor routines that individual bees might exhibit during foraging. Some bees may collect both types of nutrients, even in a single foraging bout. It would be interesting to explore whether pollen foraging on one flower species facilitates or hinders nectar foraging on another flower species depending on similarities of floral cues and required motor routines for different flower types.

### Context other than foraging

Could our results be extended to contexts beyond foraging on flowers? The underlying mechanism that governs transfer and interference effects in Osgood’s model is the generalization of similar stimuli and responses. As generalization is a universal characteristic that can be found in various animals (Lewis & Kamil, 2006), our framework has potential to be extended to other animals and other activities, offering a broader range of applications. For example, what butterflies had learned in an oviposition selection context interfered with learning stimuli in a foraging context – specifically, individuals showed a bias toward stimuli that had already been learned (Weiss & Papaj, 2003). Using our framework, one could predict a butterfly’s behaviour by assessing the degree of similarity between plants used for egg-laying and foraging, as well as the degree of similarity between egg-laying and foraging behaviours. Moreover, the framework can be extended to other learning contexts investigated in the transfer and interference literature, such as navigation (Hughes & O’Brien, 2001; Lewis & Kamil, 2006; Roberts, 1981), food caching and retrieval (Lewis & Kamil, 2006), and discrimination tasks (Dukas, 1995; Worden et al., 2005).

### Conclusion

Previous studies about interference and transfer effects have focused either on the effect of stimulus similarity or the effect of response similarity on performance of learning multiple stimulus – response associations (Hughes & O’Brien, 2001; Lewis & Kamil, 2006; Roberts, 1981). As we sought to show here, this represents only a part of the overall picture. Osgood’s model provides a unified perspective on the role of similarity in transfer and interference effects in animals. In particular, the model offers insight into the possible interaction between stimulus similarity and response similarity in the expression of memory transfer and interference. Our results provided evidence of this interaction, suggesting that further exploration of such interactions is warranted not only for bees and flowers, but for any animal that must contend with a changing environment where new stimuli and new responses of any activity must be learned.

## Acknowledgments

We are grateful to Emily Jarkowski, Melody Lau, and Liana Aguayo for their assistance in maintaining the bee colonies used in this study. We thank Chinmay Joshi for helpful comments on the manuscript.

## Data Availability

Data and code used in this study are available at https://osf.io/kzmxw/?view_only=23b86d193f7545f1b036dce6b3d6c9b2

